# K-mer clustering algorithm using a MapReduce framework: application to the parallelization of the Inchworm module of Trinity

**DOI:** 10.1101/149948

**Authors:** Chang Sik Kim, Martyn D. Winn, Vipin Sachdeva, Kirk E. Jordan

**Affiliations:** The Hartree Centre, STFC Daresbury Laboratory, Warrington, WA4 4AD, UK; Computational Science Center, IBM T.J. Watson Research, Cambridge, MA, USA

**Author notes:** Present addresses: Cancer Research UK Manchester Institute, The University of Manchester, Manchester, M20 4BX, UK. Present addresses: Silicon Therapeutics, 300 A Street, Boston MA, USA.

**Keywords:** MapReduce, *de novo* sequence assembly, RNA-Seq, Trinity

## Abstract

**Background:** *De novo* transcriptome assembly is an important technique for understanding gene expression in non-model organisms. Many de novo assemblers using the de Bruijn graph of a set of the RNA sequences rely on in-memory representation of this graph. However, current methods analyse the complete set of read-derived k-mer sequence at once, resulting in the need for computer hardware with large shared memory.

**Results:** We introduce a novel approach that clusters k-mers as the first step. The clusters correspond to small sets of gene products, which can be processed quickly to give candidate transcripts. We implement the clustering step using the MapReduce approach for parallelising the analysis of large datasets, which enables the use of compute clusters. The computational task is distributed across the compute system, and no specialised hardware is required. Using this approach, we have re-implemented the Inchworm module from the widely used Trinity pipeline, and tested the method in the context of the full Trinity pipeline. Validation tests on a range of real datasets show large reductions in the runtime and per-node memory requirements, when making use of a compute cluster.

**Conclusions:** Our study shows that MapReduce-based clustering has great potential for distributing challenging sequencing problems, without loss of accuracy. Although we have focussed on the Trinity package, we propose that such clustering is a useful initial step for other assembly pipelines.

## Background

Quantifying the expression of genes under different conditions is fundamental to understanding the behaviour and response of organisms to internal and external stimuli. With the arrival of Next Generation massively parallel sequencing technologies, the ability to monitor gene expression has been transformed [1, 2], Direct sequencing of mRNA from expressed genes (RNA-Seq) is now feasible, and has several advantages over microarray technology [3]. Most notably, it removes the need to have *a priori* knowledge of the transcribed regions, so that novel genes can be identified, or novel variants of known genes. This has led to a rapid increase in the number of studies looking at gene expression in non-model organisms. RNA-Seq is also increasingly used to study non-coding RNAs, such as microRNAs [4], lincRNAs [5], and circRNAs [6] which play various regulatory roles.

Nevertheless, it is widely recognised that the improvement in sequencing technology has shifted the bottleneck to down-stream data analysis. In the case of RNA-Seq, sequencing can be complicated by the presence of contaminant RNA, paralogous genes, and especially for higher organisms the prevalence of alternative splicing [7, 8]. Paired-end sequencing and strand-specific sequencing can help to resolve sequencing ambiguities, but must be included explicitly in the data analysis. Finally, and as we address in this study, the sheer size of datasets can cause practical problems in sequence assembly. In particular, the computational complexity limits the ability to try multiple methods or multiple parameter choices, in order to optimise the quality of the results obtained.

Initial approaches to the high throughput analysis of transcriptome sequence data were based on the alignment of RNA-Seq reads to reference genomes [9-14], Such approaches are limited by the availability of suitable reference genomes, and by the structural alterations that can be detected, particularly when input reads are relatively short Subsequently, *de novo* genome assemblers were adapted to the analysis of transcriptome data in the absence of a reference, by postprocessing draft contigs to identify transcripts. Examples of transcriptome assemblers based on genome assemblers include Oases [15] and *postprocess* [16] based on Velvet [17], TransABySS [18] based on ABySS [19], and SOAPdenovo-Trans [20] based on SOAPdenovo [21], In contrast, the Trinity [22] pipeline which we consider below was developed specifically for *de novo* transcriptome assembly. More recent examples hybridizing previous *de novo* assembly algorithms include Bridger [23] based on Trinity [22] and SOAPdenovo-Trans [20], BinPacker [24] based on Bridger [23] and bin-packing strategy [25], and DRAP [26] based on Trinity [22] and Oases [15].

Most *de novo* transcriptome assembly methods are based on *de Bruijn* graphs of k-mers, where a k-mer is a sub-sequence of an input read with k base calls. For a chosen value of k, the assembler creates a k-mer graph, where the set of nodes correspond to all unique k-mers present in the input reads, and the edges represent “suffix-to-prefix” overlaps between k-mers. Most *de novo* transcriptome assembly algorithms store all unique k-mers from the input reads in shared memory, in order to facilitate edge detection and graph construction, and this can lead to extremely large RAM usage [27], For example Velvet, as used by Oases, starts by creating two large hashmap tables in memory storing the information for all k-mers. TransABySS/ABySS is one of only a few parallel algorithms, which starts by distributing k-mers onto multiple compute nodes with a simple hash function. The Trinity pipeline consists of three independent software modules; *Inchworm, Chrysalis* and *Butterfly. Inchworm* initially creates a large hashmap table to store all unique k-mers from the input RNA-seq reads, and then it selects k-mers from the hashmap to construct linear contigs using a greedy k-mer extension approach. In our previous study [28], we confirmed that the *Inchworm* module of Trinity requires relatively high physical memory usage.

The memory requirements of these packages increase for larger and more complex transcriptomes, which generate larger numbers of k-mers and hence larger graphs, and can exceed the computational resources available. One strategy that is commonly used is to normalize the read data [29]. Redundant reads are removed from regions with high sequencing coverage, while reads are retained in regions of low coverage. In this way, up to 90% of input reads can be removed, which in turn leads to the elimination of a large fraction of erroneous k-mers associated with these reads [29]. While this is believed to work well, it introduces an additional processing step, which can in itself require large memory.

The fundamental task of *de novo* transcriptome assembly (in contrast to genome assembly) is to separate the full sequence data into many disjoint sets. Each set corresponds to a collection of gene variants sharing k-mers due to alternative splicing or gene duplication. In other words, a transcriptome can be represented as multiple distinct *de Bruijn* graphs (Fig. 1), each of which contains several paths corresponding to alternative gene products. Intuitively, *de novo* transcriptome assembly could be performed for every connected sub-graph separately. In the case of genome-guided transcriptome assembly, generation of sub-graphs is directed by the reference genome.

**Figure 1.**
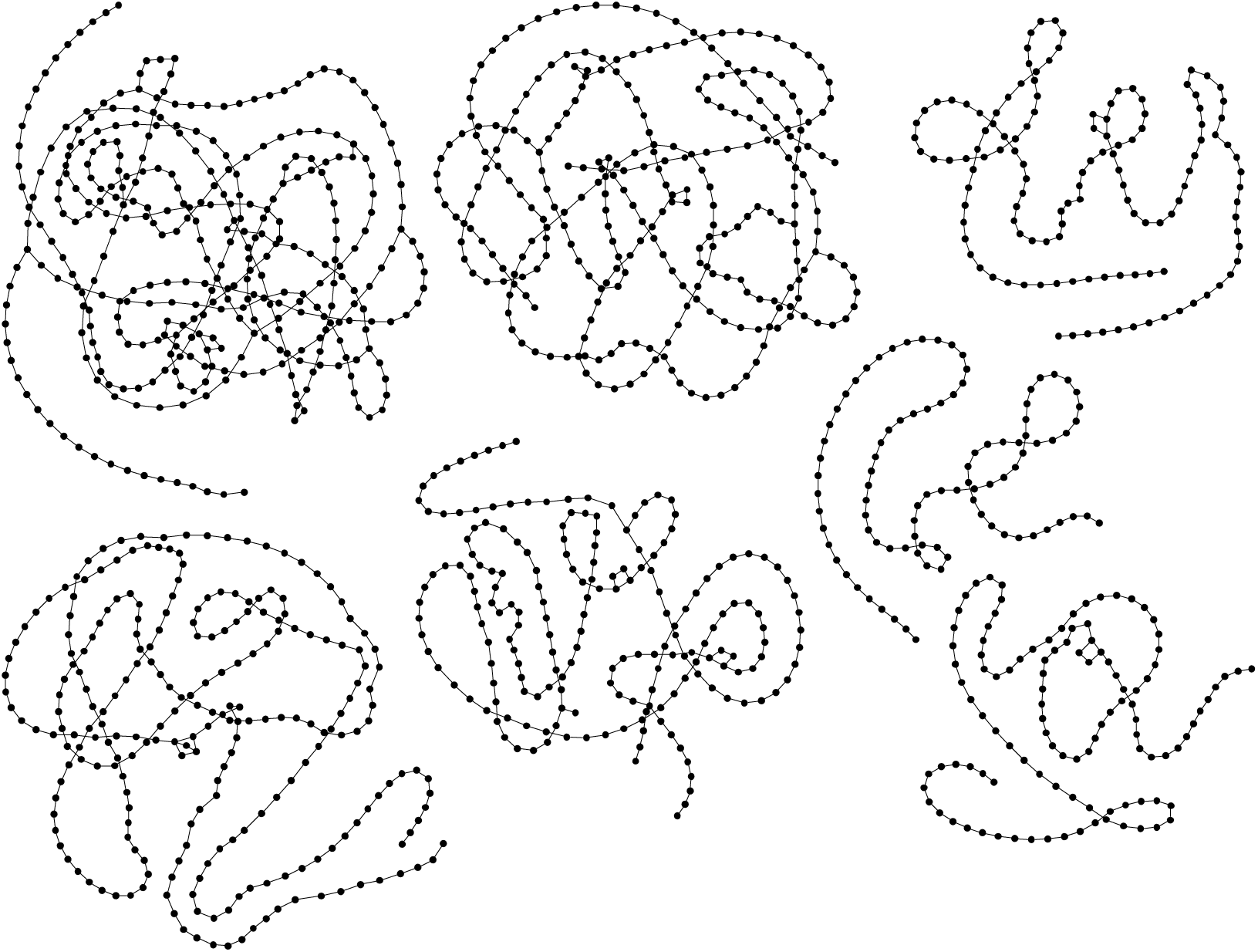
A few selected de Bruijn graphs of transcripts from whitefly RNA-Seq data. Each node represents one of the unique k-mers present in the input reads, and the edges represent *suffix-to-prefix* overlap between k-mers. Examples of branching and looping are visible (data source: http://evomics.org/learning/genomics/trinity).

In the absence of such a method for *de novo* assembly, however, most assemblers [15, 20, 22] work with all unique k-mers obtained from the input reads, resulting in the requirement for a large amount of available memory.

In this work, we present a reference-free method for generating connected sub-graphs from datasets of RNA-Seq reads. We employ the MapReduce formulation [30] for distributing the analysis of large datasets over many compute nodes. The MapReduce approach was popularized by Google for handling massively distributed queries, but has since been applied in a wide range of domains, including genome analysis [31-33]. A typical MapReduce implementation is based on *map()* and *reduce()* operations that work on a local subset of the data, but the power of the approach comes from an intermediate step called *shuffle()* or *collate()* which is responsible for re-distributing the data across the compute nodes. In the context of transcriptome assembly, the MapReduce approach distributes the sequence data over the available nodes, thus reducing the per-node memory requirement. The iterative application of *map()*, *collate()* and *reduce()* steps leads to clustering of the k-mers, such that the desired subgraphs are each physically located on a single compute node.

While distributing the sequence data across nodes of a compute cluster should lead to faster runtimes and reduced per-node memory requirements, this must be balanced against the cost of inter-node communication and transfer of data. We make use of an established MapReduce software library [34] that handles communication via the Message Passing Interface (MPI) protocol. Using this library, we have developed software that can cluster k-mers, and then launch multiple *Inchworm* jobs for the resulting sub-graphs. The procedure can be linked with the rest of the Trinity pipeline, for selected components of which we have also developed an MPI-based parallelisation [28], so that the entire assembly workflow can be run on a commodity cluster. Use of the MapReduce-MPI software library [34] means that specialised MapReduce installations such as Hadoop are not required. The only requirement is an MPI library, which is omnipresent on high performance computing platforms.

## Methods

### MapReduce-MPI library

The MapReduce [30] programming paradigm consists of two core operations, namely a “map” operation followed by “reduce” operation. These are highly parallel operations working on distributed data, which wrap around an intermediate data-shuffling operation that requires inter-processor communication. The basic data structures for MapReduce operations are key/value (KV) pairs, and key/multivalue (KMV) pairs that consist of a unique key and a set of associated values. There are many implementations of the MapReduce idea, see for examples [35, 36]. In the MapReduce-MPI library [34], which we utilise here, KV and KMV pairs are stored within MapReduce objects, and user defined algorithms consist of operations on these objects.

A typical algorithm using the MapReduce-MPI library is built upon three basic functions operating on MapReduce objects, namely *map()*, *collate()* and *reduce().* In *map()*, KV pairs are generated by reading data from files or processing existing KV pairs to create new ones. The *collate()* operation extracts unique keys and maps all the values associated with these keys to create KMV pairs. The *reduce()* operation processes KMV pairs to produce new KV pairs as input to the following steps of the algorithm. In a parallel environment, the *map()* and *reduce()* operations work on local data, while the *collate()* operation builds KMV pairs using values stored on all processors. Since KV pairs with the same key could be located on many different processors, there is a choice about where to store the resulting KMV pair. In the MapReduce-MPI library, each KMV pair is distributed onto a processor by hashing its key into a 32-bit value whose remainder modulo the number of processors is the owning processor rank.

The MapReduce-MPI library allows user-defined functions to be invoked for *map()* or *reduce()* operations, while the *collate()* operation and the general housekeeping of MapReduce objects are handled automatically. The *map()* and *reduce()* operations are called via pointers to functions supplied by the application program. Each user-defined function is invoked multiple times as a callback for each KV or KMV pair that is processed.

*Out-of-core* processing is an important feature of the MapReduce-MPI library, and is initiated when KV or KMV pairs owned by a processor do not fit in the physical memory. When this happens, each processor writes one or more temporary files to disk and reads the data back in when required. Specifically, a *pagesize* is defined by the user, which is the maximum size of MapReduce objects that can be held in memory and used in MapReduce operations. This allows the MapReduce-library to handle data objects larger than the available memory, at the expense of additional I/O to disk, and we give examples later.

### Finding Connected Components

A connected component of an undirected graph is a sub-graph where any two nodes are connected by a path of edges. A transcriptome can be represented as a k-mer graph with multiple *connected components,* where ideally the number of sub-graphs equals the number of genes (Fig. 1). The identification of connected components can be done using a depth-first search [37], Starting from a seed node, the procedure searches for the entire connected component by repeatedly looping through neighbour nodes, and creates new paths between nodes as extensions of pre-existing paths.

The algorithm starts with the assignment of unique “zone” IDs to each graph node stored in a MapReduce object. In each iteration, the size of a zone may increase by one layer of its neighbours. As zone IDs between two nodes conflict by sharing edge, a winner is chosen and the losing nodes are then merged into the winning zone. When the final iteration is reached, all nodes in a connected component will have been assigned to the same zone, and the MapReduce object will contain the zone assignments for all fully connected components. More details of the algorithm and its implementation in the MapReduce-MPI library are given in [38]. For the current application, we need to define the nodes and edges of the full (disconnected) graph to be analysed, which we do in the next subsection.

### MapReduce-Inchworm

We have implemented a multi-step procedure for clustering k-mers as the initial stages of transcriptome assembly in Trinity [22] (see Fig 2). In the first step, input sequence reads are decomposed into a list of unique k-mers, together with their abundances, as a single MapReduce cycle (Algorithm 1 in Supplementary Methods). In the second step, edges representing k-1 overlaps between k-mers are extracted in a single MapReduce operation (Algorithm 2 in Supplementary Methods). This pre-collection of edge information is an important feature of our algorithm. The third step filters out edges where a k-mer node has multiple candidates in the 3’ or 5’ directions, and is introduced to make the later *Inchworm* runs more robust (Algorithm 3 in Supplementary Methods). Inchworm builds contigs by extending a seed k-mer using the overlapping k-mer with the highest abundance, and extension continues until no more overlapping k-mers exist in the dataset. Our filtering step makes sure that the edge or edges with the highest abundance are kept in the cluster, and so available to Inchworm, while others are removed. Without this filtering operation, the subsequent step tends to produce k-mer clusters with highly diverse sizes, and leads to load balancing issues for high performance computing clusters. Having prepared the k-mer and k-mer overlap data, the fourth step (Algorithm 4 in Supplementary Methods) performs the k-mer clustering by finding connected components, as described above. The steps are described in detail in the Supplementary Methods 1.1.

**Figure 2.**
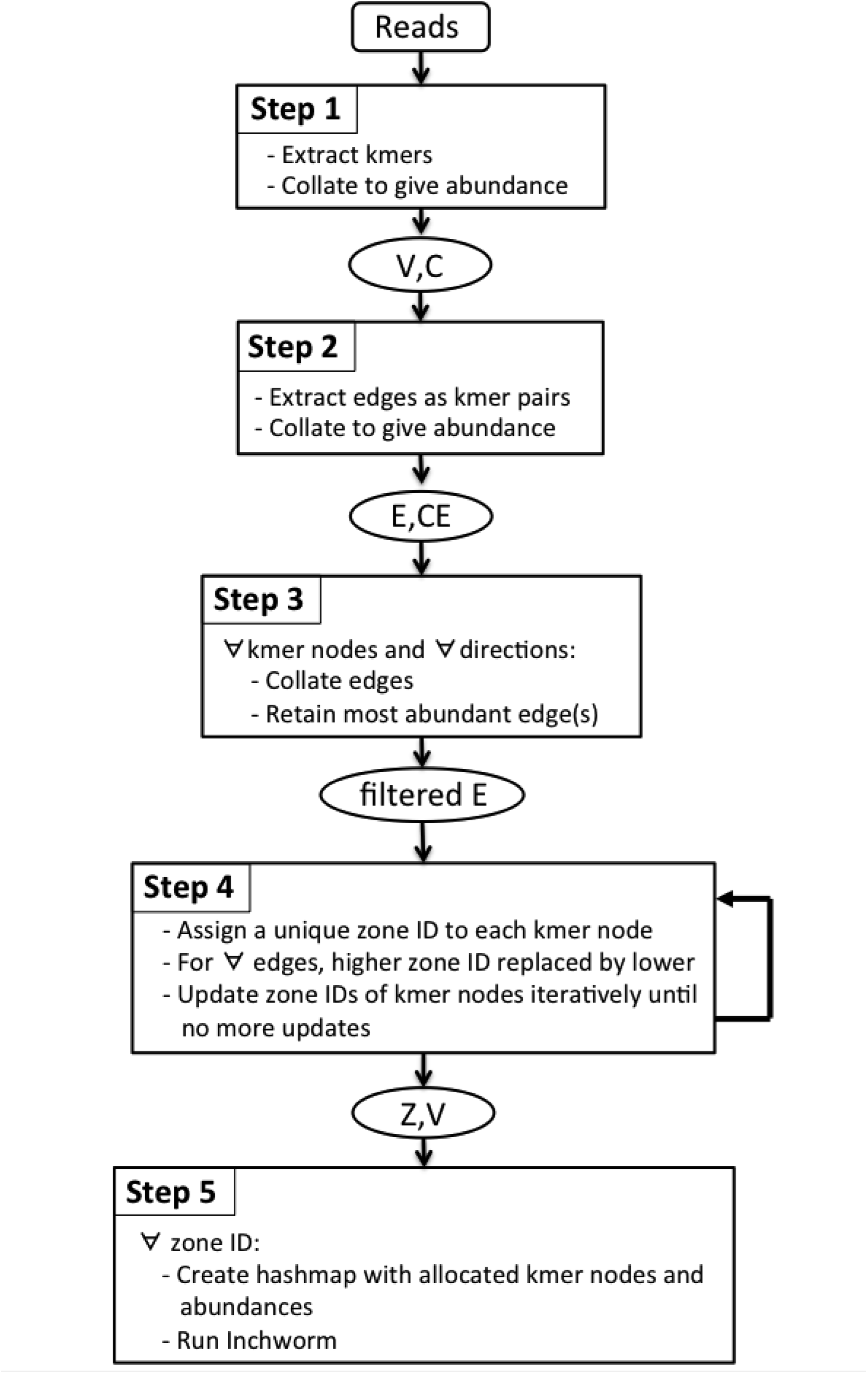
Workflow summarising the MapReduce-Inchworm algorithm. The steps are described in the main text and the Supplementary Methods. In this figure, *V* represents k-mer nodes with abundances *C*, *E* represents edges with abundances *CE*, and *Z* represents zone IDs.

The original C++ code of *Inchworm* for constructing contigs is implemented as step 5 of the algorithm, and is executed as a callback function by each set of clustered k-mers (Algorithm 5 in Supplementary Methods). The input consists of two MapReduce objects, the zone assignment of k-mers from the previous step and the list of k-mers with their abundance values. These two input objects are concatenated into a single MapReduce object, followed by a *collate()* operation using k-mers as key. This creates KMV pairs with the k-mer as key and the pair of zone ID and abundance value as the multivalue. The following *reduce()* operation creates new KV pairs, this time with the zone ID as key and the corresponding pair of k-mer and abundance as the value. Another *collate()* operation with zone ID as key produces KMV pairs with each zone ID linked to a list of k-mer/abundant value pairs.

The final *reduce()* operation creates a *hash_map* table for each zone ID, i.e. for each cluster. This table has the k-mers *V_i_* as keys and the abundance *C_i_* as values. This *hash_map* table is an input to the *Inchworm* module, which constructs contigs for that cluster. The final *collate()* operation evenly distributes the k-mer clusters across the allocated nodes of the computer. Each compute node will then run multiple *Inchworm* jobs, according to the number of k-mer clusters residing on that compute node. The resulting files of Inchworm contigs can be merged for input to Chrysalis.

## Results

This section presents our evaluation of MapReduce-Inchworm, in comparison to the original Inchworm. The primary aim of our work is to circumvent the high-memory requirements of the original Inchworm, while a secondary aim is to reduce the runtime required. It is vital, of course, that performance improvements do not lead to loss of accuracy, and so we begin by presenting a detailed characterization of the transcripts generated by the Trinity pipeline when MapReduce-Inchworm is used to generate the initial contigs. Next, we present performance results in terms of runtime and scalability, followed by results for the physical memory usage of MapReduce-Inchworm. Finally, we present a performance comparison using RNA-Seq datasets from several different organisms.

The datasets and computing resources used in our evaluations are listed in Table 1. The results presented here for MapReduce-Inchworm were obtained on an IBM iDataplex-Nextscale cluster, consisting of nodes with 2 x 12-core Intel Xeon processors and 64GB of RAM. For the original version of Inchworm, the code is necessarily run on a single node, albeit with multiple processors. For the mouse dataset, a single node of the iDataplex-Nextscale cluster was used. For the larger sugarbeet dataset, jobs were run on a high-memory (256GB) node of a slightly older iDataplex cluster. For the most complex transcriptome, the wheat dataset, ScaleMP software (http://www.scalemp.com/) was used to create a virtual symmetric multiprocessing node on the iDataplex cluster to meet the high memory requirement of the original Inchworm.

**Table 1:** RNA-Seq datasets and computing resources used for each RNA-Seq data.

| organism | # of reads | # of unique k-mers | Computing resource |  | data source |
| --- | --- | --- | --- | --- | --- |
|  |  |  | MR-Inchworm | Original Inchworm |  |
| mouse | 105,290,476 | 746,811,557 | iDataplex-nextscale | iDataplex-nextscale: single node (64GB mem) | [22] |
| sugarbeet | 129,832,549 | 2,213,519,875 | iDataplex-nextscale | iDataplex: single node (256GB mem) | unpublished data |
| wheat | 1,468,701,119 | 5,775,799,648 | iDataplex-nextscale | iDataplex: single vSMP node (4TB mem) created by ScaleMP | unpublished data |

All datasets are pair-end datasets, in which only mouse dataset is strand-specific. iDataplex-nextscale cluster is known as *BlueWonder-NextScale*, consisting of 360 nodes each with 2x12 core Intel Xeon processors (E5-2697v2 2.7GHz) and 64GB RAM making total 8,640 cores in total. iDataplex cluster is known as "BlueWonder", consisting of 512 nodes each with 2x 8 core Intel SandyBridge processors (2.6 Ghz) making 8,192 cores in total. Original Inchworm with sugarbeet dataset was run using a single iDataplex node with 256GB memory. Original Inchworm with wheat dataset was run using a single vSMP node with 4Tb memory created by ScalewMP software (http://www.scalemp.com) on iDataplex. ScaleMP creates a virtual symmetric multiprocessing (vSMP) node for shared memory by aggregating multiple compute nodes.

### Accuracy Assessment

To evaluate the accuracy of the MapReduce procedure, we compared the final transcripts generated by the Trinity pipeline when either the MapReduce-Inchworm or the original Inchworm is used. We focus on the final transcripts since these are the biologically relevant objects, while the intermediate contigs from each version of Inchworm can be quite different We performed these tests using a mouse RNA-Seq dataset consisting of 105M pair-end reads taken from [22], To generate additional datasets, we used the *rsem-simulate-reads* program from RSEM [39, 40] to simulate RNA-Seq read data based on parameters learned from the real dataset. The simulation was done in 3 steps as follows. First, we ran Trinity (using the original Inchworm) on the downloaded set of reads to produce 80,867 transcripts. These transcripts act as the set of *reference transcripts* for our trials. Second, RSEM was executed using the mouse RNA-Seq data together with the *reference transcripts* to obtain parameters for simulation of RNA-Seq reads. Third, RNA-Seq read data was simulated by executing *rsem-simulate-reads* with the *reference transcripts* and parameters from the previous RSEM run. Three simulated datasets were generated consisting of 100M, 150M and 200M pair-end reads, compared to the original experimental dataset with 105M pair-end reads, and contain approximately 5% background reads.

We ran both versions of *Inchworm* on the three simulated datasets to produce Fasta-formatted files of Inchworm contigs. The remainder of the Trinity pipeline was run from these Inchworm contigs, producing two sets of transcripts derived from MapReduce-Inchworm and the original Inchworm. The REF-EVAL module from DETONATE [41] was used to assess both sets against the “reference transcripts”, giving assembly recall and precision scores for each version of the transcriptome. Initially, all significant local alignments between assembled transcripts and reference transcripts are found using BLAT [42], At the contig level, REF-EVAL counts the number of transcripts that align with at least a predefined level of accuracy in a one-to-one mapping. We varied the required level of accuracy to get a range of statistics. At the nucleotide level, it counts the number of correctly assembled nucleotides without requiring “one-to-one” mapping; that is, it takes partially assembled transcripts into account as true positives. Recall is defined as the fraction of reference transcripts that are correctly recovered by an assembly. Precision is defined as the fraction of assembled transcripts that correctly recover a reference transcript.

We also evaluated the two quantities N1 and N2, as given by the analysis script *FL_trans_analysis_pipeline.pl* distributed with the Trinity software. This tool looks at the alignment of reconstructed transcripts onto the set of reference transcripts. If at least 99% of a reconstructed transcript is aligned to the reference, and the aligned sections have at least 99% identity, then it is considered a full-length match. The focus is on the quality of the reconstructed transcript, rather than recovery of the reference transcripts (cf. REF-EVAL above). The N1 statistic represents the total number of assembled transcripts that give full-length matches to the reference. The N2 statistic represents the number of assembled transcripts that align to multiple reference transcripts, and are thus *fused* transcripts.

The results (Table 2) show that Trinity run with MapReduce-Inchworm gives consistently higher values for Recall, Precision and N1 for the three simulated datasets. The number of fused transcripts, given by N2, is also lower. Thus, parallelisation of the initial step in the Trinity pipeline actually leads to a slight increase in assembly accuracy. In fact, the improvement of Recall and Precision at the nucleotide level is only marginal, and the absolute values are close to 1.0, indicating that both versions of Inchworm lead to transcripts that are highly similar to the reference transcripts. Recall and Precision at the contig level are lower, roughly in line with the N1 values, indicating small differences in the transcripts that lead to some reference transcripts not being fully recovered or matched. In this case, the MapReduce-Inchworm leads to a more significant improvement.

**Table 2:**
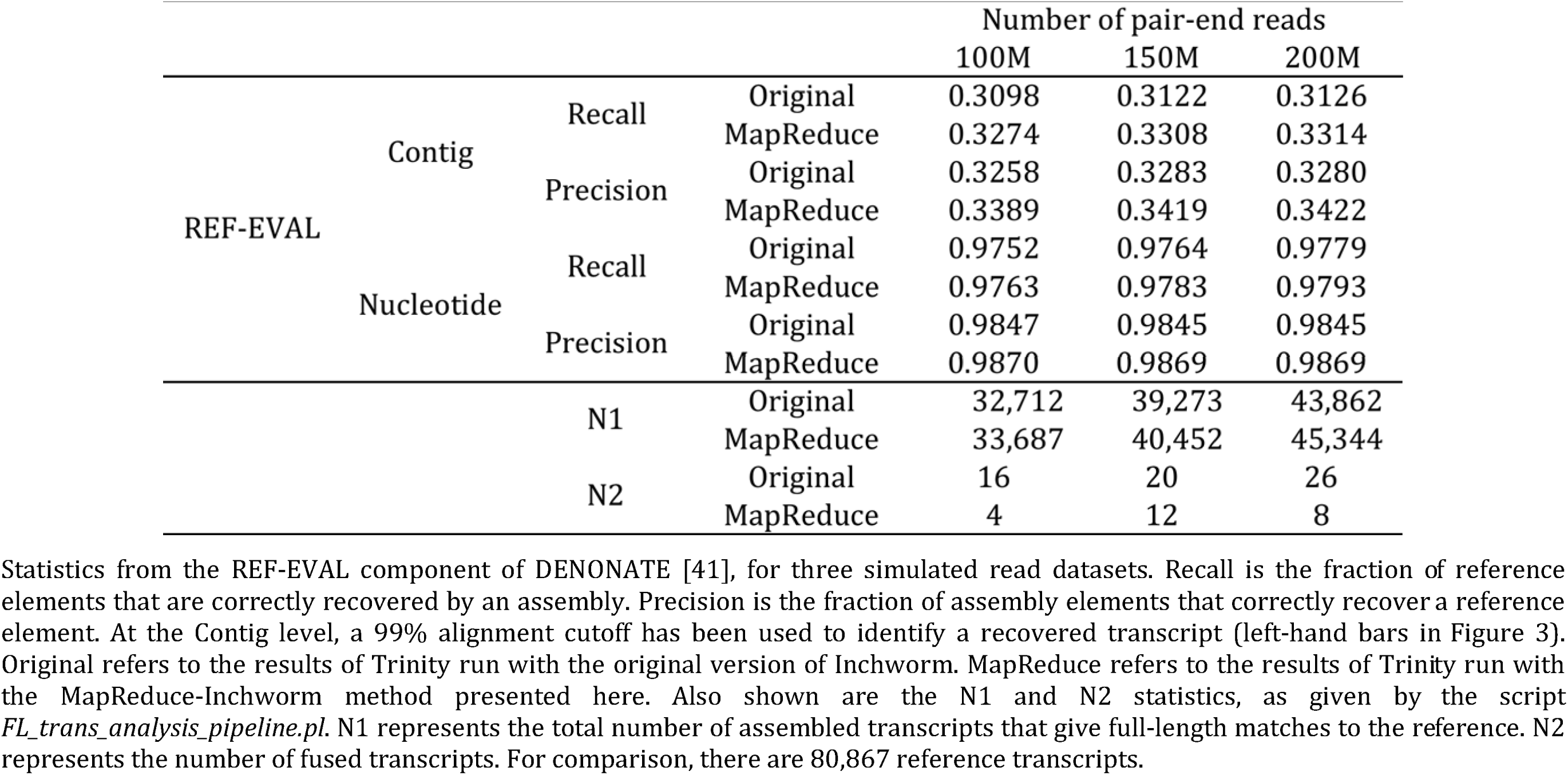
Accuracy assessment of MapReduce-Inchworm compared to the original Inchworm using three simulated read datasets for mouse RNA-Seq

Fig. 3(a) and 3(c) show the variation of the Recall and Precision statistics at the contig level, as a function of required alignment accuracy, for the simulated dataset with 100M reads. If the cutoff is reduced from 99% to 90%, so that transcripts align with high but not complete overlap, then most of the reference transcripts can be recovered from the simulated dataset. Although the absolute numbers are similar, MapReduce-Inchworm gives higher values of Recall and Precision for all cutoffs.

**Figure 3.**
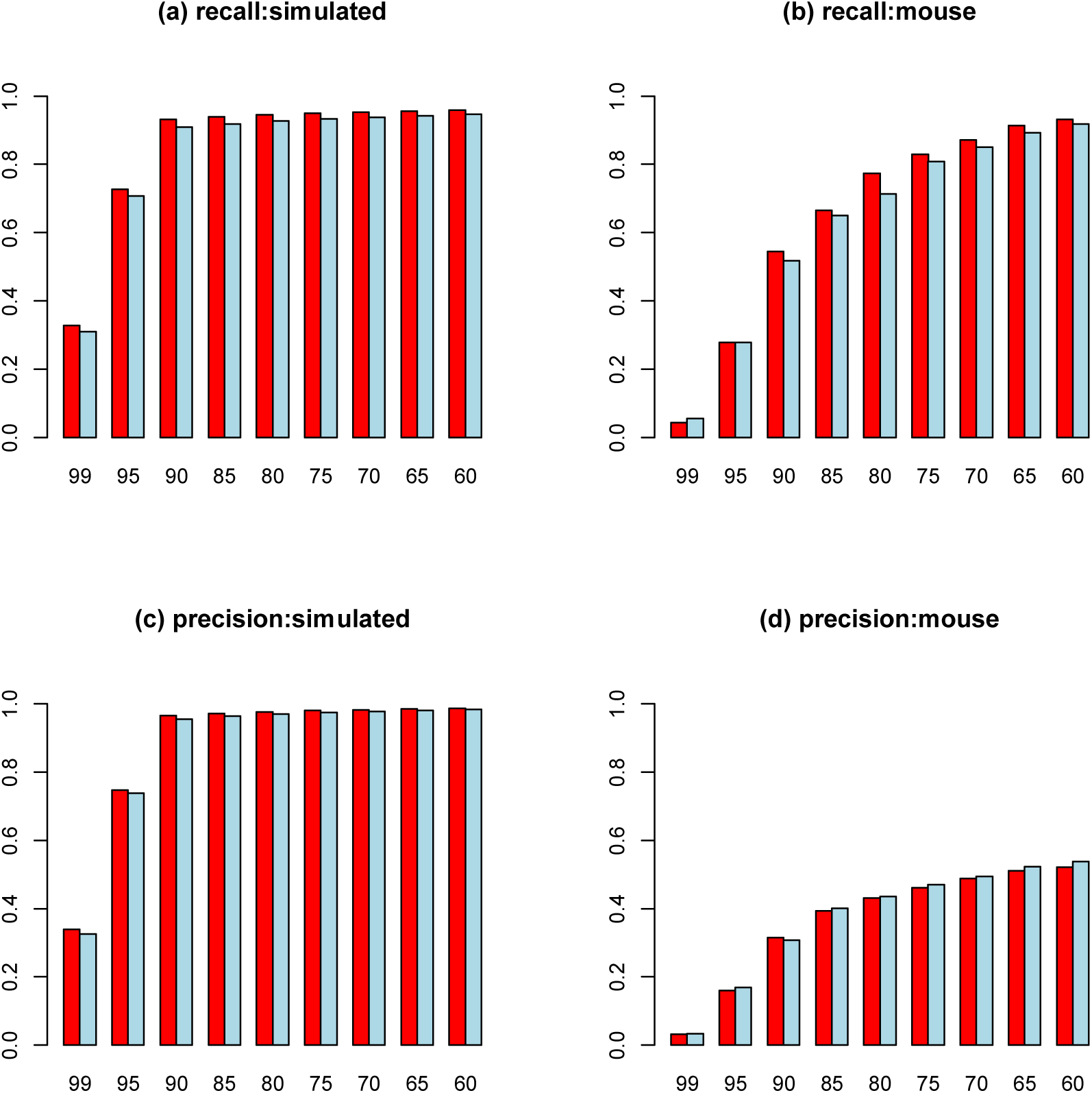
Assessment of the reconstruction accuracy of MapReduce-Inchworm (red bars) compared to the original Inchworm program (light blue bars), as given by the REF-EVAL tool of DETONATE [41]. Plots (a) and (c) give results for the dataset of 100M simulated pair-end reads (see main text), while plots (b) and (d) give the corresponding results for the original mouse RNA-Seq dataset of experimental reads. Plots (a) and (b) show the Recall statistic, which is the fraction of reference transcripts that are correctly recovered by an assembly. Plots (c) and (d) show the Precision statistic, which is the fraction of assembled transcripts that correctly recover a reference transcript Recovery of a reference transcript by a particular assembly is measured at the “contig” level, which requires almost complete alignment in a one-to-one mapping between the assembly and the reference. Each plot is given as a function of the alignment cutoff used to identify a recovered transcript.

With the simulated data, we are testing the ability of Trinity to recover the transcripts from which the simulated reads were generated. As a further test, we used REF-EVAL to compare the transcript sets that we generate to a mouse transcriptome downloaded from the UCSC genome-browser database. We used the CruzDB programmatic interface [43] to obtain a set of 22,403 coding transcripts. Statistics for the two sets of transcripts are given Table 3, and the similarity between them is quantified in Table 4. Specifically, we compared transcripts generated from the downloaded set of 105M pair-end reads using Trinity run with MapReduce-Inchworm and the original Inchworm.

**Table 3:**
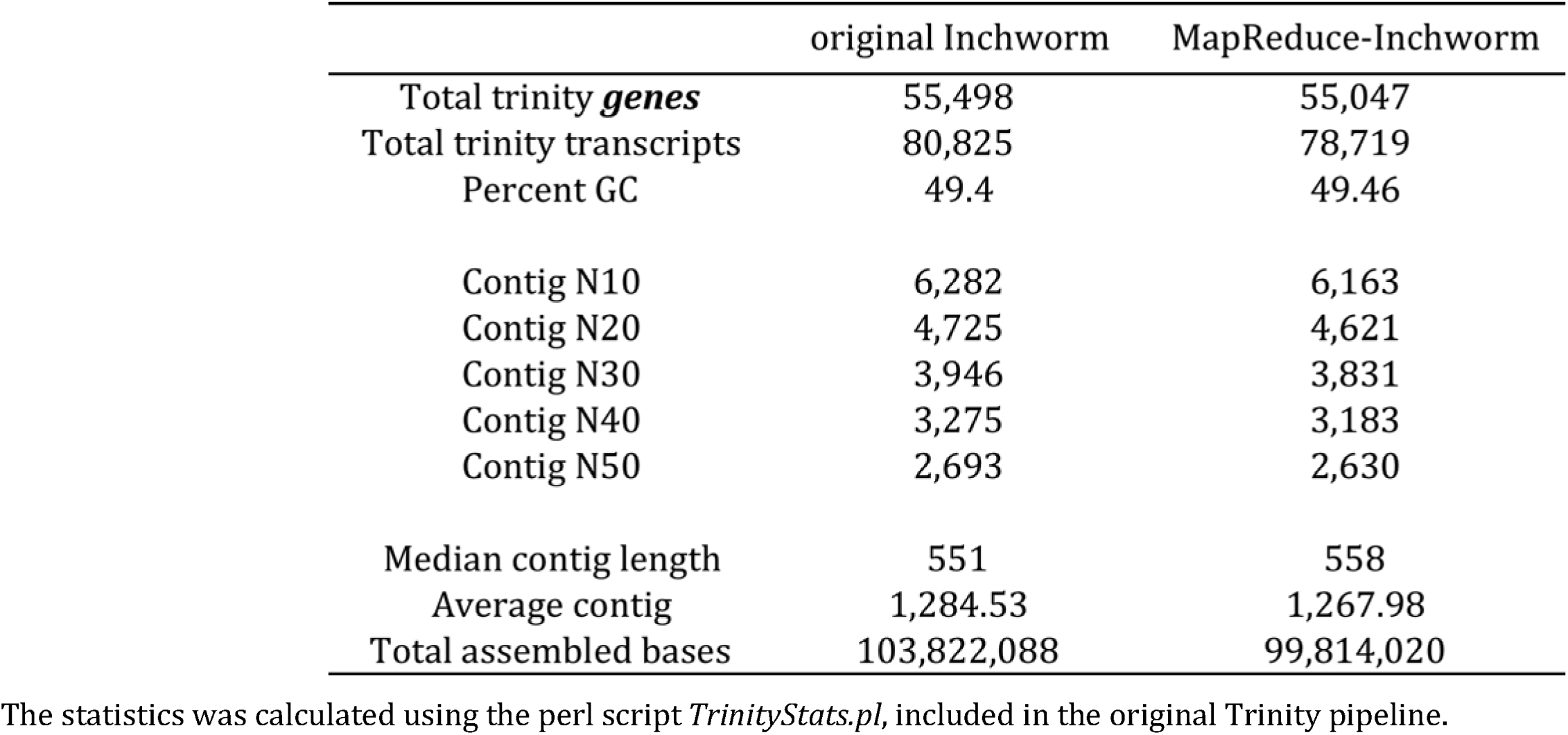
Basic statistics of Trinity transcripts using original Inchworm and MapReduce-Inchworm using the mouse RNA-seqdata [22],

|  | original Inchworm | MapReduce-Inchworm |
| --- | --- | --- |
| Total trinity *genes* | 55,498 | 55,047 |
| Total trinity transcripts | 80,825 | 78,719 |
| Percent GC | 49.4 | 49.46 |
| Contig N10 | 6,282 | 6,163 |
| Contig N20 | 4,725 | 4,621 |
| Contig N30 | 3,946 | 3,831 |
| Contig N40 | 3,275 | 3,183 |
| Contig N50 | 2,693 | 2,630 |
| Median contig length | 551 | 558 |
| Average contig | 1,284.53 | 1,267.98 |
| Total assembled bases | 103,822,088 | 99,814,020 |

The statistics was calculated using the perl script *TrinityStats.pl*, included in the original Trinity pipeline.

**Table 4:** The number of similar Trinity transcripts between original Inchworm and MapReduce-Inchworm using the mouse RNA-seq data [22],

| cutoff for transcript similarity (%) | number of similar transcripts |
| --- | --- |
| 100 | 47,816 |
| 99 | 57,926 |
| 95 | 64,109 |
| 90 | 67,178 |
| 85 | 69,002 |
| 80 | 70,398 |
| 75 | 71,390 |
| 70 | 72,285 |

Two sets of transcripts from original Inchworm and MapReduce-Inchworm were compared using BLAT [42]; Transcripts from original Inchworm was used as *target* and transcripts from MapReduce-Inchworm was used as *query* for input parameters to BLAT. The perl script *blat\_top\_hit\_extractor.pl*, included in Trinity pipeline, was used to extract the most top hit for each transcript in *query* against *target*. The first column refers to the cutoff of transcript similarity, which was quantified using two similarity score defined as follows: 1) $1 - (\text{query\_sequence\_size} - \text{number\_of\_matching\_bases}) / \text{query\_sequence\_size}$ 2) $1 - (\text{target\_sequence\_size} - \text{number\_of\_matching\_bases}) / \text{target\_sequence\_size}$ . If these two similarity scores between two transcripts from both methods were greater than or equal to the cutoff value, those were considered as similar transcripts. The second column refers to the number of similar transcripts between original and MapReduce-Inchworm according to the cutoff value. Note the total number of transcripts from both methods can be found in Table 4.

Results for contig-level Recall and Precision are shown in Fig. 3(b) and 3(d), as a function of the required alignment accuracy. The Recall is generally lower than for the simulated datasets, as the read data used probably doesn’t have the coverage to fully explain the UCSC transcriptome. Nevertheless, the Recall does approach 1.0 when the required accuracy is relaxed. The Precision also improves as the required alignment accuracy is relaxed, but remains less than 0.5 reflecting the fact that some of the read data used derives from transcripts not included in the UCSC set of coding transcripts. In the context of the current study, it is reassuring to see that again the MapReduce-Inchworm approach gives slightly improved statistics in most cases, compared to the original Inchworm (Table 5).

**Table 5:** Comparison of mouse transcripts assembled using MapReduce-Inchworm or the original Inchworm with a reference mouse transcriptome.

|  | alignment cutoff | MapReduce | Original |
| --- | --- | --- | --- |
| Number of reference transcripts recovered | 95 | 6,230 | 6,231 |
|  | 90 | 12,198 | 11,600 |
|  | 85 | 14,912 | 14,546 |
|  | 80 | 17,340 | 15,995 |
| Number of transcripts that map to reference | 95 | 12,627 | 13,601 |
|  | 90 | 24,816 | 24,770 |
|  | 85 | 30,923 | 32,397 |
|  | 80 | 33,966 | 35,208 |

We believe that the reason for the slightly improved accuracy of MapReduce-Inchworm is the inclusion of additional edge information, which is obtained in step 2 of the procedure from pairs of k-mers appearing consecutively in input reads (see Methods). With this edge information, MapReduce-Inchworm clusters k-mers into multiple groups, each of which should contain k-mers from same gene. Inchworm contigs are constructed within each cluster, and the output is implicitly guided by the input reads via this initial segregation. On the other hand, the original Inchworm uses all unique k-mers extracted from the input reads, and the construction of Inchworm contigs is done without any additional supporting information from the input reads. Both methods produce a similar total number of Inchworm contigs (data not shown), but there are clearly differences in the resulting transcripts.

### Runtime Improvement

Fig. 4 shows the scaling of the MapReduce-Inchworm runtime with increasing number of compute nodes, for the experimental mouse dataset. Plots are displayed for different choices of the *pagesize* parameter, which determines the physical memory usage (see the *MapReduce-MPl library* section in Methods for a detailed explanation). For each plot, the runtime of the original *Inchworm* (5431 seconds) is displayed as a dashed line for comparison. The number of compute nodes was varied from 32 to 192, with each node running a single MPI process, while the *pagesize* was varied from 1 GB to 4 GB. The runtimes obtained using all 192 compute nodes are 1093, 1067, 1034, and 1034 seconds for the four choices of *pagesize,* corresponding to a speed-up by a factor of about 5 compared to the original Inchworm.

**Figure 4.**
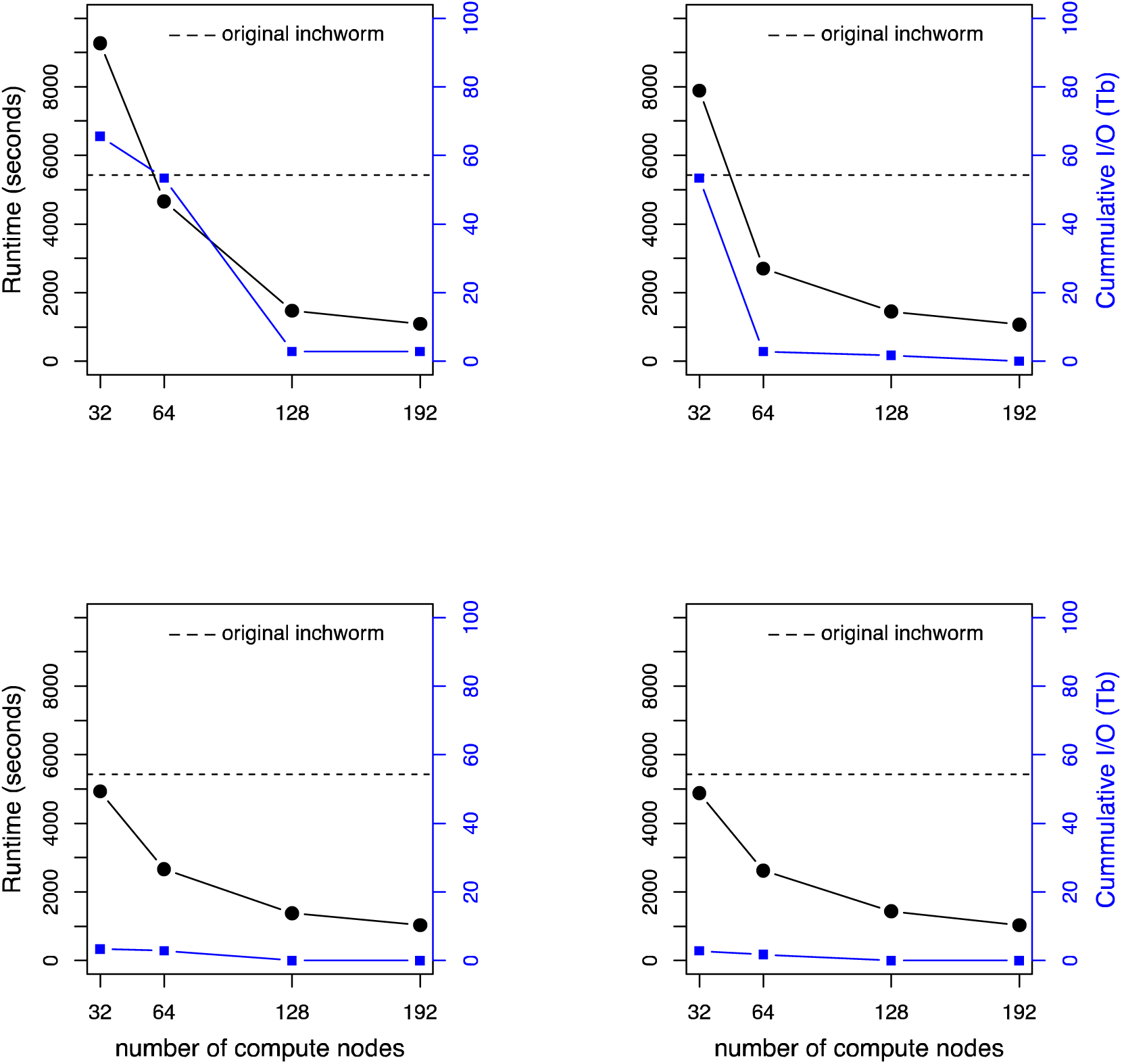
Scaling of the runtime of MapReduce-Inchworm (black lines, left-hand axis) as a function of the number of compute nodes used, for the experimental mouse dataset (see Table 1). The runtime is for the MapReduce-Inchworm step only, and does not include the remainder of the Trinity pipeline. The runtime of the corresponding serial Inchworm is shown as a horizontal dashed line. Results are shown with *pagesize* set to (a) 1 GB, (b) 2 GB, (c) 3 GB and (d) 4 GB. The cumulative I/O to disk, due to out-of-core processing, is also shown (blue line, right-hand axis).

There are also speed-ups for smaller numbers of compute nodes, except for the cases of 32 nodes with a *pagesize* parameter of 1 or 2 GB. In these cases, the memory requirements exceed the chosen *pagesize* leading to significant “out-of-core” processing (see Methods). The cumulative file I/O (Tb) is also plotted in Fig. 4, which confirms the significant paging to disk in these cases. Thus, the *pagesize* setting should be large enough (within the constraints of the available physical memory) or the number of nodes large enough (in order to distribute the memory requirements), otherwise there is an adverse effect on the runtime.

We stratified the runtime in terms of the major steps in both versions of *Inchworm,* as shown in Fig. 5. The original *Inchworm* consists of 3 principal steps: 1) *jellyfish,* 2) *parsing k-mers,* and 3) *inchworm contig construction.* The first step involves counting the occurrence of every unique k-mer in the set of input reads using the program Jellyfish [44], and writing the output to a disk file. In the second step, *Inchworm* reads the output file back into physical memory by storing each k-mer and its count into a hashmap table as a key-value pair. In the final step, the algorithm creates draft contigs using the hashmap table of unique k-mers. We divide the MapReduce-Inchworm algorithm into an initial *splitting input reads* step, followed by the five steps described in Methods. The initial step consists of evenly splitting the input file of reads into multiple files, according to the number of allocated compute nodes. Each file is then read into a compute node in preparation for subsequent steps.

**Figure 5.**
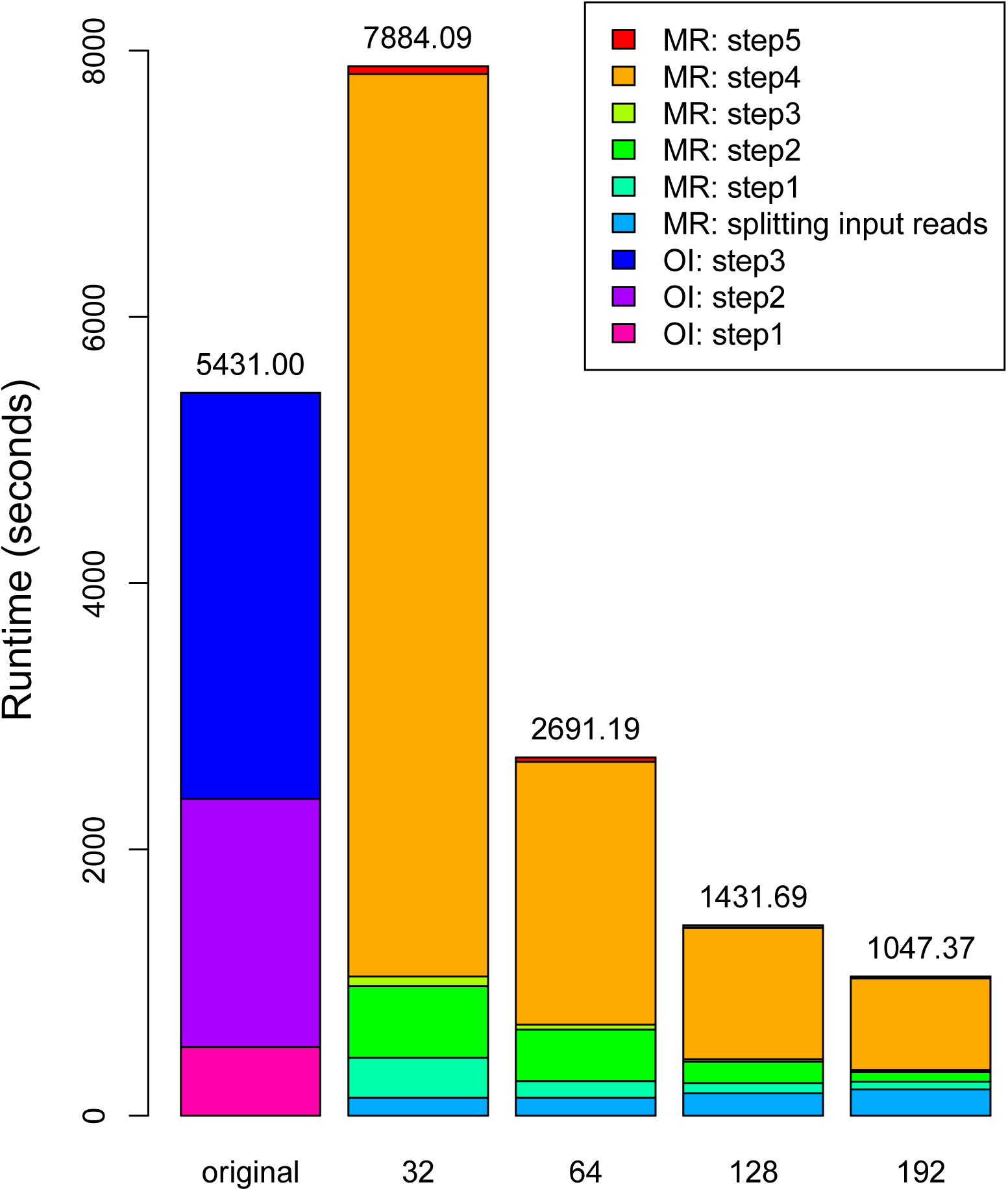
Stratification of the runtime in terms of individual steps within both versions of Inchworm, for the experimental mouse dataset (see Table 1). 0I represents Original Inchworm; MR represents MapReduce-Inchworm. On the X-axis, *original* represents the original version of Inchworm, while 32-192 represent the numbers of compute nodes allocated for MapReduce-Inchworm. The original Inchworm is divided into three steps: step1 corresponds to *Jellyfish* [44]; step 2 corresponds to *parsing kmers;* and step3 corresponds to *Inchworm contig construction.* MapReduce-Inchworm is divided into six steps: an initial step for *splitting input reads* and steps 1-5. The initial step splits an input file (containing the RNA-Seq reads) into multiple files according to the number of allocated compute nodes. Steps 1 to 5 of the main algorithm are described in detail in Methods. Results are given with *pagesize* assigned to 2GB, cf Figure 3(b).

Fig. 5 shows that the first two steps of the original Inchworm, which could be categorised as k-mer preparation steps, take a significant fraction of the total runtime. These steps are equivalent to the *splitting input reads* and step 1 of MapReduce-Inchworm. The latter steps are however much quicker because they avoid storing k-mers on disc. The remaining runtime of the original Inchworm involves construction of contigs. In the MapReduce-Inchworm implementation, this is done individually for each cluster, and is very fast (MR: step 5 in Fig. 5). The bulk of the runtime for MapReduce-Inchworm is taken by the clustering algorithm (MR: step 4 in Fig. 5), and this scales well with the number of nodes used. As mentioned above, super-linear scaling is achieved in going from 32 nodes to 64 nodes because of the reduction in out-of-core processing, while going from 64 to 128 nodes gives a speedup of 1.9, and from 64 to 192 nodes a speedup of 2.6.

### Physical Memory Requirement

The main objective of our work is to remove the need for large shared memory, by distributing the overall memory requirement over multiple computer nodes. With the ability to do that, the per-node memory requirement can always be reduced by adding more compute nodes, albeit with the expense of increased inter-node communication. The physical memory available on each node is controlled by the *pagesize* parameter in the underlying MapReduce-MPI library. In this section, we look at the memory requirements of MapReduce-Inchworm, as a function of the number of compute nodes and the *pagesize* parameter.

Firstly, we assessed the memory requirements as a function of the number of allocated compute nodes, using the mouse dataset (see Table 1). Within the 5 main steps of MapReduce-Inchworm, we collected the number of KV/KMV pairs generated from each of the three basic MapReduce functions: *map, collate,* or *reduce.* These values were converted into data object sizes in GB, and averaged over all compute nodes. Fig. 6(a) shows that the data size per compute node, and hence the memory requirement, decreases with increasing number of nodes, as expected. The values for step 4 are also averaged over the iterations of the k-mer clustering algorithm, of which there are 47 for the mouse dataset. The figure shows clearly that step 2 of the MapReduce algorithm, which extracts edges from the input read data, is the most memory demanding.

**Figure 6.**
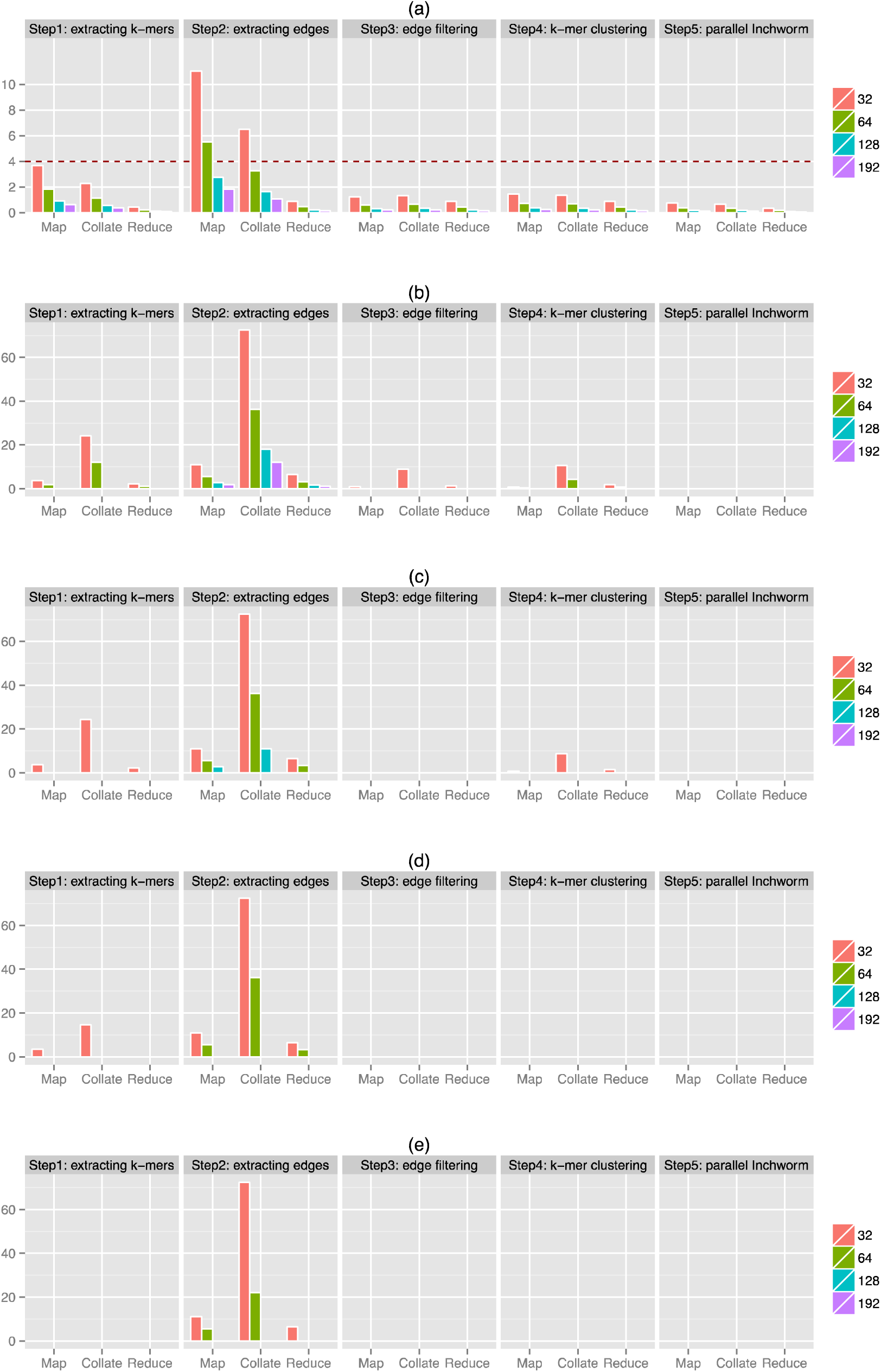
Data objects created by MapReduce-Inchworm for the experimental mouse dataset (see Table 1). (a) The size of data objects generated by each MapReduce function on each compute node, as calculated from the number of KV/KMV pairs involved. The data sizes for each MapReduce function (*map()*, *collate()*, and *reduce()*) were averaged over nodes and iterations within each of the 5 main steps of MapReduce-Inchworm. (b-e) The corresponding cumulative I/O to disk, due to out-of-core processing, per compute node. Results are shown with *pagesize* set to: (b) 1 GB, (c) 2 GB, (d) 3 GB and (e) 4 GB. For all graphs, the Y-axis gives the data size in GB.

The values in Fig. 6(a) give an estimate of the per-node memory requirements of MapReduce-Inchworm. When these exceed the physical memory allocated according to the *pagesize* parameter, then pages of data are written as temporary files on disk. Paging for each of the steps is shown in Fig. 6(b)-(e) for four choices of the *pagesize* parameter. For example, the data sizes of KV pairs obtained from the *map* operation of step 2 are 11.0~GB, 5.5~GB, 2.75~GB and 1.83~GB when run on 32, 64, 128 and 192 nodes respectively (see Fig. 6(a)). For a small *pagesize* of 1 GB (Fig. 6(b)), there is always some out-of-core processing. Increasing the *pagesize* to 2 GB (Fig. 6(c)) means that, in the case of 192 compute nodes, the KV pairs can fit in memory and there is no paging. Increasing the *pagesize* to 3 GB (Fig. 6(d)) means that out-of-core processing is eliminated when using 128 or 192 compute nodes. However, there is always paging for the smaller compute clusters considered, even for the largest tested *pagesize* of 4 GB. Out-of-core processing in the *map* operation of step 2 initiates *out-of-core* processing in the following *collate* step, because the KV pairs written to disk by the *map* operation need to be read back for the *collate* step. Fig. 6(b)-(e) shows that file I/O occurs for the *collate* operation of step 2 whenever it is present for the *map* operation. It also shows that total I/O is substantially higher for the *collate* operation. This arises because the core *collate* algorithm needs to cycle through the KV pairs multiple times, as it finds matches and builds up the KMV objects, necessitating multiple reading and writing of pages to disk. Thus, step 2 of the MapReduce algorithm is particularly sensitive to the choice of the *pagesize* parameter. On the other hand, Fig. 5 shows that the runtime is dominated by step 4, and so the performance hit caused by paging in step 2 is perhaps not so important.

The underlying MapReduce-MPI library tends to evenly distribute the KMV pairs produced by *collate* operations by hashing each of its key for assignment onto available MPI-processors. In the present application, where the number of KMV pairs is much larger than the number of compute nodes, this is expected to lead to good load balancing. In fact, the minimum and maximum values for data size over all compute nodes are indistinguishable from the average values shown in Fig. 6(a).

### Performance comparison for RNA-Seq datasets from complex organisms

We next tested our approach on some more challenging RNA-Seq datasets obtained from sugarbeet and wheat samples, see Table 1. The memory requirement of the original *Inchworm* depends on the transcriptome complexity and is expected to roughly correlate with the number of unique k-mers from the input reads. Fig. 7(a) shows that the mouse, sugarbeet and wheat datasets require 46.7GB, 141.5GB and 373.9GB of memory respectively, and these values do indeed correlate with the total number of unique k-mers listed in Table 1. In fact, the required memory for the wheat dataset exceeded the physical memory on any single node of our available compute platforms. In order to run the original *Inchworm,* we used ScaleMP software (http://www.scalemp.com/) to aggregate 32 nodes, each providing 128GB memory, to create a vSMP node with a 4TB address space.

**Figure 7.**
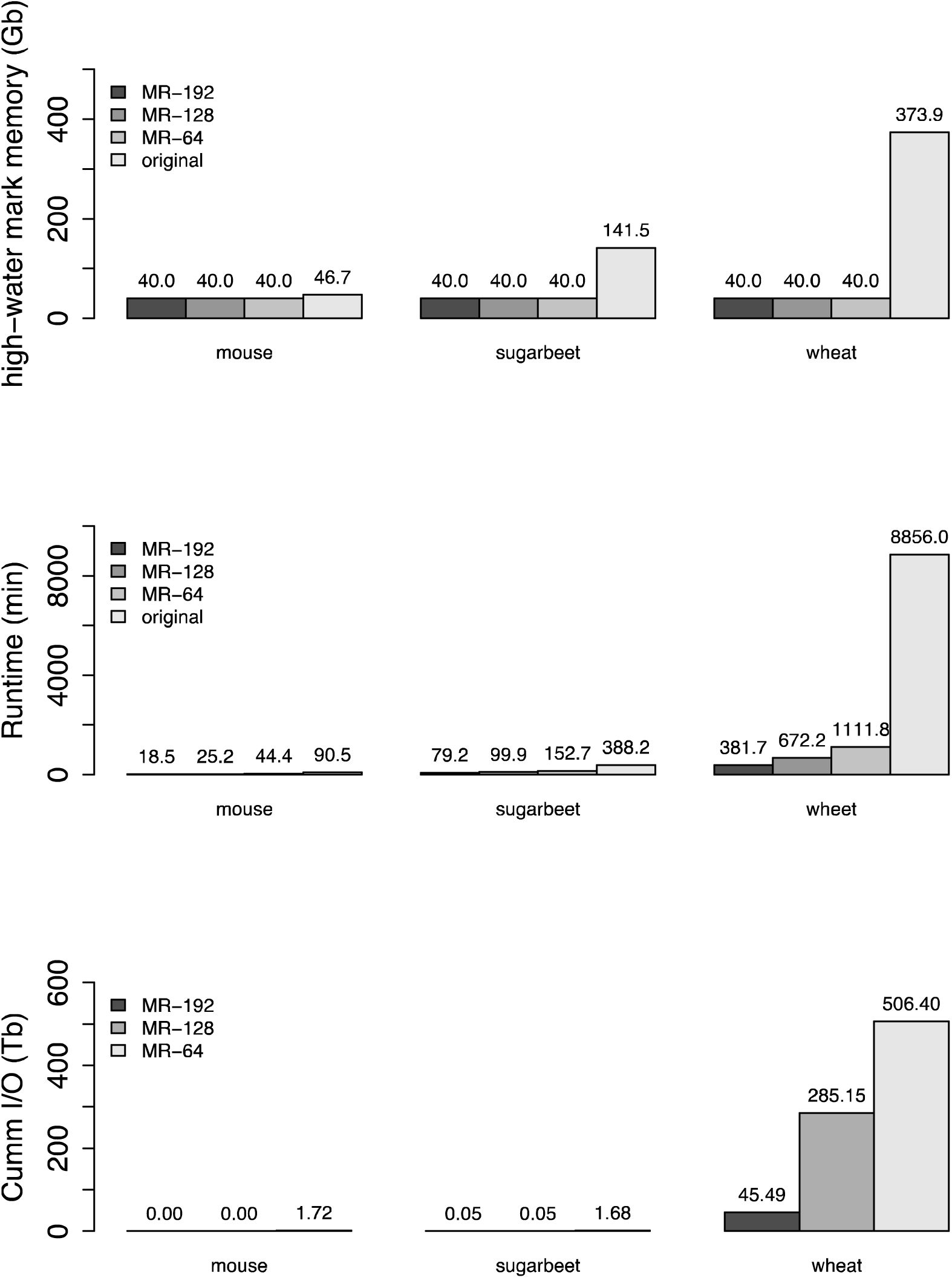
(a) High-water mark for memory usage in GB over all compute nodes, (b) runtime in minutes, and (c) cumulative I/O in TB. Results are given for the mouse, sugarbeet and wheat datasets described in Table 1, and using the computing resources listed. All jobs with MapReduce-Inchworm used a 4GB *pagesize* parameter. Bars marked *original* represent runs of the original Inchworm, MR-64 represents runs of MapReduce-Inchworm using 64 compute nodes, MR-128 represents runs using 128 compute nodes, and MR-192 represents runs using 192 compute nodes.

Fig. 7(b) shows the total runtime to produce *Inchworm* contigs for *MapReduce-Inchworm,* compared with a run using the original *Inchworm.* With 64 compute nodes available, the *MapReduce-Inchworm* procedure yields a faster runtime, and increasing the number of nodes to 128 or 192 gives further improvements. Although we have only tested a small range of node counts, the scaling of the speedup is very good, suggesting that more nodes could be used. For example, tripling the number of nodes from 64 to 192 yields speedups of 2.4, 1.9 and 2.9 for mouse, sugarbeet and wheat respectively. The original *Inchworm* is particularly slow for the wheat dataset, taking over 6 days, because of the use of ScaleMP to provide the required amount of shared memory. While ScaleMP was required to be able to process the wheat dataset at all, its software-based aggregation of memory clearly incurs a significant overhead. Such problems are avoided in the distributed memory implementation of *MapReduce-Inchworm.*

Fig. 7(c) shows the cummulative I/O for the runs using *MapReduce-lnchworm,* reflecting the *out-of-core* processing of MapReduce-MPI. The wheat dataset, with 1.5~billion input reads and 5.8~billion unique k-mers, is particularly large and the MapReduce objects do not fit into the aggregate physical memory. Even with 192 nodes, there is 45TB of I/O over the course of the run. This could be reduced by allocating further compute nodes, or by using higher memory nodes that could accommodate a larger pagesize. For the smaller sugarbeet dataset, the situation is much better, but there is nevertheless still a small amount of paging to disk.

In conclusion, the *MapReduce-lnchworm* procedure scales well to large and complex datasets. Increasing the number of compute nodes leads to a reduction in runtime, and reduced paging to disk as the per-node memory requirements are lowered. In particular, *MapReduce-lnchworm* allowed us to process the large wheat dataset in less than a day, while the original *Inchworm* required an advanced platform solution to run at all.

## Discussion

In this study, we enabled the parallelization of the *Inchworm* module of Trinity by using a MapReduce-based approach to pre-cluster the k-mers obtained from the input reads. An instance of *Inchworm* is run on each k-mer cluster, yielding a set of contigs per cluster. Contigs from all clusters are pooled together and passed to the *Chrysalis* module for re-clustering according to the original Trinity scheme. The *Inchworm* module of Trinity is known to be the most memory-intensive step [28], and is often a barrier to processing large or complex RNA-Seq datasets. In our scheme, the computational load is passed to the pre-clustering step, where the well-established MapReduce procedure allows the load to be distributed over a commodity compute cluster. Our approach is distinct from other recent developments, which seek to MPI-parallelise *Inchworm* itself (Brian Haas, personal communication).

Pell et al. [45] have introduced a Bloom filter which provides memory efficient storage of kmer graphs. Chikhi and Rizk [46] added an additional data structure holding the false positives which might arise through trial kmer extensions, as well as a marking structure holding complex nodes for use in graph traversal. In contrast, we store an explicit list of kmer nodes and edges in a set of MapReduce objects. There is no reduction in the total memory required, but rather we focus on the ability to distribute these MapReduce objects over a large number of nodes to reduce the per node memory requirement. Note that by storing edges explicitly, we do not make trial extensions from kmer nodes, which can lead to false edges in the Bloom filter method.

We expect that there should be a correspondence between the k-mer clusters we generate, and the contig clusters (*components*) produced by *Chrysalis,* in that they both relate to a set of gene products. It may be that further efficiency gains can be achieved by merging these steps, but we have not investigated this possibility in the current work, adopting instead a conservative approach. If, for example, reads from a transcribed gene yield two k-mer clusters, and hence two sets of *Inchworm* contigs, then the *Chrysalis* module should in principle find *welds* between them, and recover the correct graph.

The assessment of the assembly accuracy using simulated and experimental RNA-Seq datasets shows that our parallelized *Inchworm* provides the final transcripts from the Trinity pipeline with marginally more accuracy compared to the original inchworm. The difference in accuracy comes from the utilization of additional edge information in MapReduce-Inchworm, which clusters k-mers guided by edge objects which link pairs of k-mers appearing consecutively in input reads. On the other hand, the original *Inchworm* constructs contigs directly from the set of all k-mers. Contigs are extended by searching for appropriately overlapping k-mers, rather than using pre-calculated edges. This pre-collection of edge information is feasible in our approach because of the distributed nature of the algorithm.

We have presented performance results for a range of experimental read datasets. The total runtime required to produce the complete set of *Inchworm* contigs could be reduced below that required for the original *Inchworm,* provided a moderate number of compute nodes are available. It may be debatable whether this is necessary for small datasets, such as the mouse dataset included here, but there are clearly significant gains for larger and more complex datasets (Fig. 7). More importantly, the memory requirement on each node can be reduced by distributing the job over sufficient nodes. In this way, commodity compute clusters can be used, and there is no need for high memory nodes or specialised solutions for aggregating node memory into a single address space.

## Conclusion

The results of this study indicate that the MapReduce framework has great potential for processing high throughput sequencing datasets more efficiently. The proposed approach could be applied as a pre-processing step for other *de novo* transcriptome assemblers, by implementing the chosen assembly code as a callback function in the final *reduce()* step, as we have done for *Inchworm* in the current study. Specifically, we plan to investigate the parallelization of Oases [15] and SOAPdenovo-Trans [20] via this approach. It is also worth exploring the feasibility of the pre-clustering approach for *de novo* metagenome and metatranscriptome assembly, which is more complex due to the presence of multiple genomes or transcriptomes from different species. For example, the *de novo* metagenome assembler [47] starts by partitioning the *de Bruijn* graph into isolated components corresponding to different species. Then for each component, it reconstructs the slight variants of the genomes of subspecies within the same species using multiple sequence alignments. A similar approach has been developed for *de novo* metatransciptome assembly [48]. Our proposed approach could be adapted to these pipelines to provide a memory-efficient method for the initial partitioning.

In conclusion, we have presented a computationally efficient method for clustering k-mers derived from RNA-Seq datasets. Applied to the Trinity pipeline, the approach avoids the large memory requirements of the original Inchworm, enabling the analysis of large datasets on commodity compute clusters. We expect that this general approach will have applications for other assembly problems.

## Declarations

### Acknowledgement

We would like to thank Brian Haas of the Broad Institute for his advice on the Trinity software and source code. We thank Keywan Hassani-Pak of Rothamsted Research, UK for providing the sugarbeet and wheat datasets, as well as many useful discussions. We thank Steve Plimpton for making the MapReduce-MPI library freely available, and for answering technical questions on its use. We also wish to acknowledge the Hartree Centre at the STFC Daresbury Laboratory, UK for providing the computational resources used for this work.

### Authors’ contributions

CSK and MDW conceived the study. CSK implemented the method. CSK and MDW designed the research. All authors analysed the data. The benchmarking and repository maintenance were done by CSK and MDW. All authors wrote, read and approved the final manuscript.

### Availability of material

The code used in this study is available at https://github.com/kimosaby2001/MR-Inchworm/.

### Consent for publication

Not applicable.

### Ethics approval and consent to participate

Not applicable.

